# FINS - A global occurrence dataset of fossil neoselachians from the Cretaceous to the Quaternary

**DOI:** 10.1101/2025.11.21.689757

**Authors:** Kristína Kocáková, Jaime A. Villafaña, Amanda Gardiner, Gregor H. Mathes, Catalina Pimiento

**Affiliations:** Department of Paleontology, University of Zurich, Zurich, Switzerland; Departamento de Ecología, Facultad de Ciencias, Universidad Católica de la Santísima Concepción; Concepción, Chile; Department of Genetics, University of Cambridge, Cambridge, United Kingdom; Chair of Physical Geography, University of Passau, Passau, Germany; Department of Biosciences, Swansea University, Swansea, United Kingdom

**Keywords:** Cretaceous, Cenozoic, fossil occurrences, Neoselachii, rays, sharks and skates

## Abstract

**Motivation:** Modern sharks, rays and skates (Neoselachii) are a diverse, globally distributed, and ecologically important group. They possess a rich, geologically extensive, and well-documented fossil record that provides a powerful basis for studying their macroevolution, paleoecology and biogeography. However, addressing paleobiological questions require quantifying the fossil record trough occurrences, i.e., records of the presence of a taxon at a unique location in space and time. Although the Paleobiology Database (PBDB) offers an invaluable source of fossil occurrences, it still does not provide a complete compilation of neoselachian data. To facilitate exploring questions in quantitative paleobiology, here we present the Fossil Neoselachian (FINS) dataset—a comprehensive, systematically curated global synthesis of fossil occurrences spanning the past 145 million years.

**Main types of variables contained:** FINS consists of fossil occurrences, each linked to a collection (location where fossils were reported from) and a source (citation of the publications from which data was obtained). Occurrence data mostly consists of taxonomic variables (e.g., classification, names and ranks), status (extinct/extant) and abundance. Collection data provide geographic (e.g., country, ocean and paleo-coordinates) and geological (e.g., formation and age) variables. All data was subjected to a meticulous curation process, which is described through taxonomic and age validation variables.

**Spatial location and grain:** Global, with fossil occurrences recovered from all continents.

**Time period and grain:** From the Cretaceous to the Quaternary (145 – 0 Ma), with temporal resolution generally corresponding to geological stages or epochs.

**Major taxa and level of measurement:** Neoselachians (superorders: Selachimorpha and Batoidea). FINS comprises 30,100 occurrences from 5,972 collections and 1,990 publications. It consists of 1,606 species, 512 genera, 88 families and 14 orders. Over 70% of occurrences are identified at the species level. Overall, FINS effectively doubles the number of neoselachian records previously available in the PBDB and substantially expands their geographic coverage, particularly in the Global South.

**Software format:** The dataset is provided an *.xlsx* spreadsheet with four tabs – Collections, Occurrences, References_Literature and References_PBDB.

## 1. Introduction

Modern sharks, rays and skates, and their close extinct relatives, form the monophyletic group Neoselachii (Cappetta, 2012; Maisey et al., 2004). The neoselachian fossil record possesses several characteristics that make it stand out amongst other fossil marine vertebrate groups. First, despite their rarely preserved cartilaginous skeletons, neoselachian teeth (and denticles) have high preservation potential due to their hard composition and their continuous growth and shedding (Cappetta, 2012; Hubbel, 1996; Wang & Cerling, 1994). As such, neoselachians are one of the most abundant vertebrate groups in the marine fossil record, and have a relatively well-preserved evolutionary history (Maisey et al., 2004). Second, the neoselachian fossil record extends deep in geological time, with the first fossils dating back to the Triassic (250 million years ago, herein Myr) and some extant genera being traceable to the Jurassic (190 Myr; Paillard et al., 2021). Third, despite high levels of interspecific variation in tooth morphology (Cappetta, 2012; Guinot et al., 2018), neoselachian teeth can be diagnostic, allowing for a well-resolved classification, including identifications to the species level (Maisey, 2012). Finally, neoselachian tooth morphology has been shown to be a good proxy of biological traits such as body size and diet, enabling ecological inferences (J. A. Cooper et al., 2023). In summary, the neoselachian fossil record is abundant, geologically extensive and biologically informative.

Due to these characteristics, neoselachian fossils have been a prominent subject of study in palaeontology since the birth of the field, as exemplified by the seminal work of Louis Agassiz in the early 19^th^ century (Agassiz, 1833). Even though many important descriptions of neoselachian fossils were made through the last 200 years, the neoselachian literature has become more prominent over the last 50 years, specifically from the 1970s. This increase in publications can be attributed to a shift from surface collection of fossils, a method which favoured larger specimens, to bulk sampling, which allows for much smaller specimens to be collected (Cappetta, 2012), significantly increasing the number of identified taxa (Underwood, 2006). A synthesis of this knowledge is, however, still lacking.

The growing number of studies on fossil neoselachian over the last five decades has opened new opportunities to address fundamental questions in paleobiology, including how clades diversify, compete and respond to global changes through deep time (Brée et al., 2022; Condamine et al., 2019; Guinot et al., 2012; Guinot & Condamine, 2023; Kriwet et al., 2009; Kriwet & Benton, 2004; Kriwet & Klug, 2008; Staggl et al., 2025; Underwood, 2006; Villafaña et al., 2023; Villafaña & Rivadeneira, 2018). The exploration of these questions, however, require quantitative approaches, with fossil occurrences (i.e., records of the presence of a taxon at a specific place and time) representing the core empirical units of analysis. The Paleobiology Database (herein PBDB; www.paleobiodb.org; Uhen et al., 2023) offers an invaluable repository of such data. However, despite the efforts of hundreds of contributors, the PBDB still does not offer a complete synthesis of neoselachian occurrences. As a result, most of recent works on neoselachian paleobiology have relied on independently compiled occurrence datasets (e.g., Guinot & Condamine, 2023; Kriwet & Benton, 2004; Villafaña et al., 2023).

Here, we present the Fossil Neoselachian (FINS) dataset, a comprehensive, systematically curated global synthesis of fossil occurrences spanning the past 145 Myr. FINS has two main objectives: (1) to synthesise existing knowledge of the neoselachian fossil record; and 2) to provide a standardized, curated and easily accessible compilation of fossil occurrences to facilitate their use in quantitative paleobiology. The dataset focuses on the Cretaceous to Quaternary interval (145–0 Ma), capturing the most recent phase of neoselachian evolutionary history that underpins their present-day diversity. This timespan encompasses several significant global events, such as the Cretaceous/Paleogene mass extinction (K/Pg), Paleocene-Eocene Thermal Maximum (PETM), and Eocene-Oligocene transition.

To build FINS, we first downloaded the existing data from PBDB. Then, we identified the records not yet included in this source, extracted them from the literature and combined them following the PBDB data structure (Fig. 1). To locate additional records, we used Shark-References (herein SR, www.shark-references.com; Pollerspöck & Straube, 2022), an online platform compiling all known literature on extant and extinct cartilaginous fishes (class: Chondrichthyes) (Fig. 1). We focused on the literature from 1970 onwards, which represent the most productive period for research on the neoselachian fossil record. SR primarily provides bibliographic metadata, with additional information including geographical region, temporal intervals and the taxa mentioned in each publication. SR also contains a continually curated and updated list of valid taxonomic names, synonyms, and taxonomic classifications, which is separate from the literature collection. In line with our first objective (i.e., synthesising current knowledge), FINS faithfully captures information from the literature, similar to PBBD. However, in line with our second objective (to provide a standardized, curated and easily accessible compilation of fossil occurrences), all records in FINS underwent a systematic curation process that involved updating taxonomic and stratigraphic information based on the most reliable and recent sources. This curation process is fully transparent and thoroughly documented, enabling users to trace each record and apply alternative criteria if desired.

**Figure 1.**
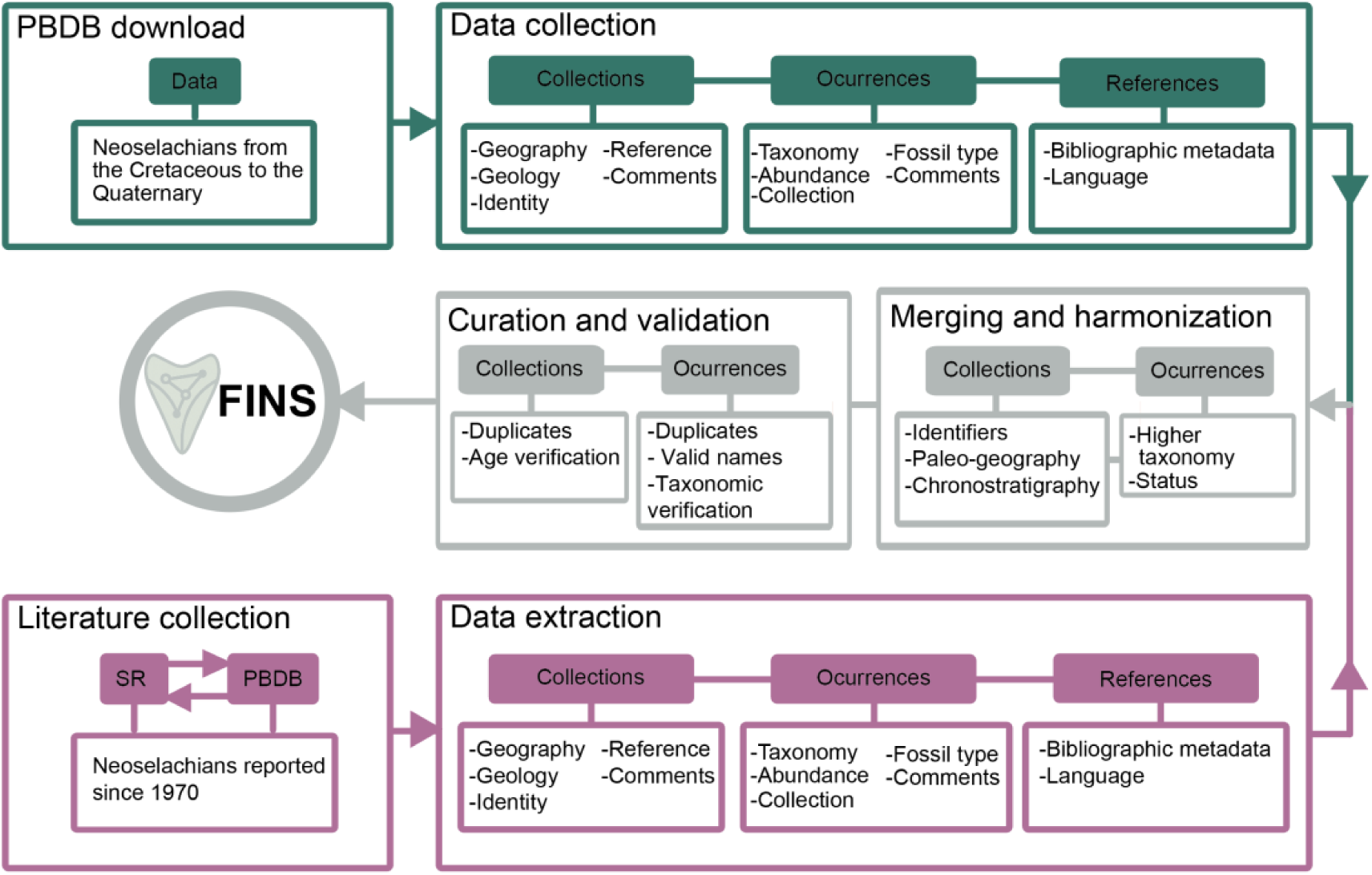
Workflow of the FINS dataset creation. Data was either gathered directly from the literature or downloaded from PBDB. Three types of data were collected: collections, occurrences and references. These data were merged enriched and harmonized into the FINS dataset and then critically evaluated. SR denotes Shark-References, PBDB denotes Paleobiology Database.

The final dataset comprises 30,100 occurrences from 5,972 collections from 1,990 publications in 16 different languages. It consists of 1,606 species, 512 genera, 88 families and 14 orders, with temporal resolution generally corresponding to geological stages or epochs. To date, FINS represents the most comprehensive global compendium of neoselachian fossil occurrences available, enabling the identification of both large-scale patterns and persistent data gaps. Notably, occurrences were recovered from all continents, substantially expanding the geographic coverage of existing PBDB data, particularly in underrepresented regions of the Global South. Neoselachian fossils are present in every epoch, with abundance peaks in the Late Cretaceous, Eocene, and Miocene, and the fewest records in the Holocene. Beyond its descriptive scope, FINS provides a robust foundation for investigating macroevolutionary patterns such as diversification and extinction dynamics, for reconstructing paleobiogeographic histories including past range shifts and provinciality, and for exploring species responses to environmental change.

## 2. Methods

### 2.1. Data structure and PBDB download

The PBDB provides global collection-based occurrence and taxonomic data for all taxonomic groups and ages. It is broadly organised in three primary data classes: occurrences, collections and bibliography (Uhen et al., 2023). An occurrence is a record of a taxon at a unique location in space and time. A collection is the location where an occurrence is reported from, with unique coordinates and age. Each occurrence and associated collection is linked to a reference of the publication from which the information was extracted from. In some cases, there can be multiple occurrences of the same taxon in a collection if they come from different publications reporting new material. Similarly, multiple publications about the locality can result in different collections, especially if they are published across different decades. Finally, a locality reporting fossils from multiple geological horizons with different ages represent different collections (one for each age). FINS was built following this data structure (collections, occurrences and references).

To build FINS, we first downloaded collection, occurrence and reference data of all Neoselachii from the Cretaceous to the Quaternary from the PBDB (last accessed June 2020, Fig. 1). This data was accumulated through the effort of numerous contributors, which we acknowledge in Table 1. We used the following criteria for our download query: taxon to include = Neoselachii; interval = Cenozoic AND Cretaceous. For the reference download, we selected the option “collection references”. We manually checked the downloaded output to ensure that only neoselachians (superorders Selachimorpha and Batoidea) were included. All records were considered, regardless of the year of publication. Collection information includes collection number, country, state, latitude, longitude, formation, member, stratigraphic scale, collection name, collection aka, early interval, late interval, minimum and maximum age, and reference. Occurrence data include collection number, occurrence number, identified name, abundance and reference. Although extinct and extant status of a taxon is provided in PBDB, we have added this information ourselves (see Methods). The reference data includes the full citation of all the publications from which data was extracted from. We added the prefix *PBDB_* to each collection and occurrence number, to differentiate them from those collected from the literature (see below).

**Table 1.** Overview of contributors who entered neoselachian occurrences to the PBDB.

| Number of contributions | Contributors |
| --- | --- |
| > 1000 | Uhen M., Clapham M., Pimiento C., Carrano M. |
| 500 - 100 | Mannion P., Glaeser W., Leckey E., Holroyd P., Alroy J., Hendy A., Vlachos E., Villafaña J., Marcot J. |
| < 100 | Layou K., Varnham G., Nurnberg S., Krause M., Shalap M., Benson R., Gagucas M., Wall P., Oreska M., Mordaunt J., Zijlstra J., Eccles L., Tennant J., Bell M., Sessa J., Novack-Gottshall P., Cleary T., Nelson M., Villier L., McMullen S., Cárdenas A., Alfaro O., Fischer V., Behrensmeyer A., Kiessling W., Kouvari M., Maguire K., O'Regan H., Peredo C., Yu E., Fara E., Head J., Seuss B., Merkel U., Furich F., Egerton V., Medeiros E., Street H., Aspromonte F., Hulbert R., Pino K., Schossleitner T., Ballen G., Allen B., Butler R., Lloyd G., Kriwet J., Saeed A., Low S., Kosnik M., Cote S., Visaggi C., Volkman K., Wertheim J., Clyde W., Schweitzer C., Kahane N., Carillo J., Coster P., McVicker C., González Tejos M., Whatley R., Chan C., Villari J., Wagner P., Valdez R., Vazquez P., Mora-Rojas L., Devicezi S., Buczek A., Dalal A., Jablonski D., Liebrecht T., Millet J., Tredwell Z., Tasimov E., Jaramillo C., Brand N., Benefield J., Alam S., Lin C., Edwards T., Araya S. |

### 2.2. Literature collection and data extraction

To locate literature that reported neoselachian occurrences not yet included in PBDB we used SR (last accessed October 2020; Fig. 1). SR is an online database listing the literature describing or mentioning chondrichthyan fossils in any language. The database contains metadata for each publication, including publication type (e.g., journal articles, conference proceedings, book chapters, specialised magazine articles and student theses), geographical and chronostratigraphic keywords, and lists of described taxa. We obtained a list of all the references in SR and contrasted it to that from PBDB. To optimise our data extraction (see below), only literature published from 1970 onwards was considered given that the last 50 years cover the vast majority of neoselachian reports (Underwood, 2006). Literature which reported Synechodontiform taxa was included in the literature list, as this order is considered to be a part of Neoselachii *sensu* Klug (2010). We excluded references reporting only Hybodontiformes or Holocephali, as these clades do not belong to the monophyletic Neoselachian group despite being closely related (Maisey, 1984; Villalobos-Segura et al., 2022).

In total, we identified 851 publications in SR that reported neoselachians but that that were not yet included in the PBDB. We carefully read each of these papers and extracted data using the PBDB structure as follows:

#### 2.2.1 Collections

For each publication considered, we searched the localities reported in the PBDB to ensure it was not already entered there. If a new collection was identified, an identifier consisting of a prefix and a number was given to it and reported in the “collection_number” column. Accordingly, all collection numbers from the literature have an *L_* prefix to differentiate them from those downloaded from PBDB (*PBDB*_ prefix). Six categories of information were collected for each collection (Fig. 1) as described below.

##### Collection identity

A name (“collection_name” column) was given to each collection based on the locality name and any alternative names used (“collection_aka” column).

##### Geographical information

We extracted country, state (or other type of local region, e.g., prefecture, municipality, province), latitude and longitude. If specific coordinates were not provided in the source literature, they were derived based on a nearby landmark (e.g., city, mountain, lake). If such derivation was done, this was noted in the “collection_comments” column.

##### Stratigraphical information

We extracted the geostratigraphic units of each collection. This information was entered in the columns “formation”, “member”, and “stratscale”, the latter reporting the stratigraphic resolution to which the units are known, e.g., bed, member, group of beds etc. Whenever this information was not provided in the publication, it was left blank.

##### Age information

Each collection has a geochronological interval (columns “early_interval” and “late_interval”; e.g. Cenozoic, Paleocene-Eocene, etc.) associated with it, which we extracted from the publications. The majority of publications report intervals based on the International Chronostratigraphic Chart (Walker & Geissman, 2022), however, some reported region-specific units (Table 2). The resolution of the chronostratigraphic unit was reported in the “time_interval_type” column (e.g., stage, epoch, sub-stage, etc.).

**Table 2.** Stratigraphic units which were reported in some publications and assigned to collections, that do not follow the International Chronostratigraphic Chart (Cohen et al., 2013). The references presented here were used to assign ages in millions of years to these collections.

| Abbreviation | Name | Reference |
| --- | --- | --- |
| ELMA (=MN) | European land mammal age (= Mammal Neogene Zones) | Agustí et al., (2001) |
| MP Zone | Mammal Paleogene zone | Taken from individual publications |
| NALMA | North American Mammal age | Alroy, (2017); Fowler, (2017) |
| NZ Stage | New Zealand Stage | Raine et al., (2015) |
| Regional stage | Stages of Central Parathetys | Kováč et al., (2018) |
| SALMA | South American Land Mammal Age | Flynn & Swisher, (1995) |

##### Reference

Each collection is associated with the reference (“reference” column) which the information was extracted from. This includes the first author and year of the publication, e.g. Carrillo-Briceño et al. 2015.

##### Fossil type

We extracted the type of fossils reported from each collection and reported it in the “fossil_type” column. Fossil type was assigned to one of the following categories: teeth, vertebrae, fin spines, dental plates, rostral spines, dermal denticles, gill rakers, skeletal remains (e.g., jaw fragments, cranial remains and cartilage), imprints, complete specimen, coprolites, capsule egg, or unspecified).

#### 2.2.2 Occurrences

Similar to collections, each occurrence was assigned a numerical identifier with an *L_* prefix to differentiate occurrence obtained directly from the literature from those obtained from the PBDB (*PBDB_* prefix, “occurrence_number” column). Each occurrence is associated with the collection from which it was identified, provided in the “collection_number” column. If occurrences were reported from a collection which was already entered in the PBDB, we reported these occurrences with an *L_* prefix and reported the collection number from PBDB in the “collection_number” column. Six categories of information were collected for each occurrence (Fig. 1).

##### Taxonomic information

The taxon name exactly as given in the literature is reported in the “identified_name” column.

##### Abundance

If the publication reported the number of fossils associated with the occurrence (e.g., number of teeth), this number was reported in the “abundance” column, otherwise this column was left blank.

##### Comments

Any noteworthy information about the occurrence is noted in the “occurrence_comments” column.

#### 2.2.3 References

Bibliographic information is reported in the columns “author”, “year”, “title”, “journal”, “DOI” and “abstract”, the last two only filled when available. If author and year were shared by multiple publications, they were differentiated by a letter (e.g., a, b, c), provided in the “identifier” column. If the publication was a thesis, the “journal” column contains the university at which the thesis was published. For the other types of literature, this column contains the name of the publication, such as book name or magazine name. The first author (“author1” column) is provided as an additional option for easier navigation. The “journal_detail” column contains additional information such as the volume and issue number. The type of publication (e.g. journal article, thesis, abstract) is specified in the “pubtype” column. The language in which the publication is written is reported in the “language” column.

### 2.3 Data merging and harmonization

The data downloaded from PBDB and the data extracted from the ∼800 publications from SR was merged, enriched and harmonized. Additional geographical and stratigraphic information was added to the collections. Similarly, chronostratigraphic, geographic and taxonomic information was linked to each occurrence. This process was done was follows.

#### 2.3.1 Collections

##### Identifiers

We assigned an identifier to each collection in the “locality_id” column. This allows users to identify instances in which a collection was reported in the literature as consisting of multiple localities of the same age. As each locality in these cases was assigned to a unique collection number, there are several collections in FINS with identical coordinates and age, but different locality identifiers (e.g., L_1164 to L_1170).

##### Number of occurrences

The number of occurrences (“n_occs” column) in each collection was recorded.

##### Geographical information

Additional geographical information added to collections included continents, latitudinal bands, paleo-coordinates and paleo-ocean. Continents were assigned based on the country column using the ‘pycountry’ package (v. 22.3.5; Theune, 2023) in Python (v. 3.7; Rossum & Drake, 2010). As Russia spans two continents, Russian collections were assigned to a continent based on their longitude, with a cut-off value based on the position of the Ural Mountain Range (i.e., 60.33°), which splits the country between Europe and Asia. Whenever there was a collection from an of open ocean locality, we manually assigned the continent to “ocean”. Latitude bands were assigned to each collection based on the “latitude” column, assigning each collection to one of the following bands: northern temperate (i.e., north of the Tropic of Cancer, 23.44°), tropical (i.e., between the Tropics of Cancer and Capricorn) and southern temperate (i.e., south of the Tropic of Capricorn, −23.44°).

Paleo-coordinates for each collection were calculated based on the modern coordinates of each collection (rounded to a single decimal point) and their age, using a global gridding R package ‘dggridR’ (v. 3.1.0) and plate tectonic models accessed via the ‘chronosphere’ (v. 0.4.1) R package (Barnes, 2018; A. T. Kocsis et al., 2024; Á. T. Kocsis & Raja, 2019; Müller et al., 2018). It should be noted that paleogeographic functions of ‘chronosphere’ were moved into the ‘rgplates’ R package, however at the time of creation of this dataset this was not the case, thus ‘chronosphere’ was used directly. The reconstruction of paleo-coordinates varied depending on the age of the collection. To account for this age-uncertainty, we created thirty replicates from a uniform distribution delimited by the minimum and maximum age for each collection. Each replicate was re-rotated to calculate its paleo-coordinates, and we used the median value as our estimate. The majority of paleo-coordinates were calculated using the Matthews 2016 model (Matthews et al., 2016). However, paleo-coordinates for some collections located near tectonic plate boundaries could not be quantified due to the more dynamic interactions, such as subduction, rifting, and transform faulting, returning missing values. For these collections, we used an alternative model (PALEOMAP; Scotese, 2016) which features an alternate set of plate boundaries. Despite this, the paleo-coordinates for seven collections remained unresolved, as they still fell near plate boundaries in the new model. Five of these could then be resolved based on the Seton 2012 model (Seton et al., 2012), and the remaining two collections via the Müller 2019 model (Müller et al., 2019). All models mentioned above were accessed through ‘chronosphere’. Paleo-oceans were then assigned to each collection based on paleo-coordinates. We first obtained the age range and locality of each paleo-ocean basin from literature (Scotese, 2014a, 2014b, 2014d, 2014c, 2017). We then assigned an ocean basin to each paleo-coordinate replicate and calculated the modal value (i.e., the ocean basin which was assigned most often across replicates), in order to propagate the age uncertainty and the resulting differences in plate reconstructions (see Data and Code availability). Even though paleo-coordinates are available in the PBDB, in order to keep consistency of the data, these values were calculated for PBDB collections in the same way as for data extracted from literature.

##### Age information

Additional chronostratigraphic information was manually added to each collection based on the “early_interval” and “late_interval” columns, following the International Chronostratigraphic Chart (ICC; version 2022/10; Cohen et al., 2013) or following the appropriate literature if alternative region-specific intervals were used in the source literature (Table 2; columns: “early_epoch”, “late_epoch”, “early_period”, “late_period”, “early_era”, and “late_era”). Region-specific stages sometimes overlap with two ICC defined epochs, for example Mangapanian, a New Zealand stage, spans both Pliocene and Pleistocene. In such cases, the epoch which shared a longer time span with the stage was assigned. The age range in millions of years (Ma) of each interval was also assigned based on the ICC or literature (columns “max_ma” and “min_ma”), unless higher-resolution ages (e.g. based on isotope dating) were provided in the publications. These instances are noted in the “collection_comments” column. Finally, the “age_range” column was calculated and assigned to each collection based on the “min_ma” and “max_ma” columns (see Data and Code availability).

##### Notes

The column “collection_comments” describes any noteworthy aspects found during the data gathering, such as the source of coordinates, other non-neoselachian taxa described, and any updates made to ages (see 2.4 Data curation and validation).

#### 2.3.2 Occurrences

##### Status

Whether or not a taxon was extinct or extant was assigned to each occurrence in the “status” column. Extant taxa were identified based on a list compiled previously (Pimiento et al., 2023). The “genus_status” column indicates whether the genus which a species belongs to is extant or extinct, even if the species reported is extinct.

##### Higher taxonomy

Higher taxonomy was assigned to each occurrence and reported in the “family”, “order”, and “superorder” columns, based on the taxonomic list form SR. Note that this step was only carried out after the curation and validation process (see 2.4.2. *Synonyms and valid names*). The higher taxonomy was therefore assigned based on the valid names (see Data and Code availability).

##### Collection information

The following columns were added to each occurrence based on the collections associated with them: “max_ma”, “min_ma”, “age_range”, “early_interval”, “late_interval”, “early_epoch”, “late_epoch”, “early_period”, “late_period”, “early_era”, “late_era”, “int_type” (interval type), “latitude”, “longitude”, “continent”, latitude_band”, “paleoocean”, “paleolat” (paleolatitude), “paleolon” (paleolongitude), and “locality_id”(see Data and Code availability).

#### 2.3.3 References

##### Collection information

Chronostratigraphic, geographic and taxonomic information associated with each reference was added. This is included in the columns “keyword_time”, “keyword_place”, “described_species”, “described_genus” and “described_fam_ord”. Taxa described for the very first time in a given publication are listed in the columns “nov_fam”, “nov_gen”, “nov_sp”. Note, that this information was only collected for literature from which we extracted data, but not for references from PBDB.

### 2.4 Data curation and validation

#### 2.4.1 Record duplicates

After the merging and enriching of the data, a check for duplicates was carried out. In the case of collections, a duplicate would be a collection with the same or very similar name (i.e., slight difference in spelling) and/or with the same coordinates, age and stratigraphy. If a duplicate was found, the PBDB version was kept, and the collection number associated with the occurrences was updated to the PBDB collection number. In some cases, duplicates were found in occurrences as well, where two occurrences representing the same taxon were reported in one collection. A part of these duplicates resulted from occurrences being taken from a thesis, abstract or a poster that was later published as a peer-reviewed article. In these cases, only the occurrences described in the peer-reviewed articles were kept. Duplicates were kept if a taxon was reported in a given locality multiple times by different authors. These duplicates were retained in the FINS dataset as they represent the true state of the literature, however they should be addressed appropriately before any analyses (see Usage below).

#### 2.4.2 Synonyms and valid names

For each occurrence, in order to assign valid names to outdated synonyms, we first manually identified and modified names with typos and with taxonomic qualifiers (e.g., sp., sp.nov., ex gr.). These modified names are reported in the “modified_identified_name” column, which was then used to manually assign a taxonomic rank (“rank” column; e.g., species, genus etc.) to each occurrence. Modified names at the species- and genus-level were then cross-checked with the taxonomy from SR which includes a list of valid names and synonyms (last accessed in March 2021; Data S1). Valid names in SR are based on Cappetta (2006, 2012) and Ginter et al. (2010) and are regularly updated by its curators. Therefore, the SR taxonomy is dynamic and changes over time. The cross-checking was carried out as follows. The taxonomic list from SR was converted into a Python dictionary of synonyms and valid names, which was then used to match the names given in the “modified_identified_name” to the appropriate values (see Data and Code availability). Whenever synonyms were identified, valid taxonomic names were reported in the “accepted_name” column. Taxonomic names that did not need updating were recorded in the “accepted_name” column as given in the “identified_name” or the “modified_identified_name” columns. Similarly, occurrences with taxonomic ranks of family and higher were reported as such in the “accepted_name” column.

#### 2.4.3 Taxonomic verification

The validity of the taxonomic information associated with each occurrence was evaluated in the “occurrence” dataset. We treated uncertain nomenclature (i.e., containing ?, cf., aff, etc.) as valid and maintained the taxonomic rank of the occurrence (i.e., species, genus, etc.). However, both the original taxonomic name provided by the publication (e.g., *Archaeolamna aff. kopingensis*) and the one deemed to be valid (e.g. *Archaeolamna kopingensis*) are accessible in the dataset in the “identified_name” and “accepted_name” columns for both transparency and usage flexibility.

For the data collected from the literature, we checked whether evidence was provided for the taxonomic assignment of each occurrence. We considered evidence to be present if: (a) the source publication included an image of the specimen; (b) a museum accession number was provided; or (c) a detailed description was provided (e.g., in a Systematic Palaeontology section). If such evidence was present in the publication, the occurrence was marked as *with_evidence* in the “evidence_validation” column, otherwise it was marked as *no_evidence* and an explanation noted in the “evidence_comments” column. For the PBDB data, the statement on whether taxonomic evidence was provided was downloaded for each reference obtained from the PBDB and entered into the “evidence_comments” column. If evidence was provided according to the PBDB, the associated occurrences were marked as *with_evidence* in the “evidence_validation” column, otherwise they were marked as *no_evidence*.

Although we assigned valid names to each occurrence based on the SR taxonomic list (see above), there were several instances in which the taxonomic name could not be verified because it was not present in the list (e.g., *Asiadontus shuvalovi*). In such cases, the occurrence was marked as *invalid* in the “taxonomy_validation” column, as *unverifiable* in the “accepted_name” column, and an explanation was noted in the “taxonomic_validation_comments” column. A few PBDB occurrences (0.22% of total PBDB occurrences) were marked as *invalid* despite having a valid name assigned, as they had been assessed in previous works and deemed unreliable (e.g., occurrences PBDB_14360-63; Pimiento et al., 2016; Pimiento & Clements, 2014).

#### 2.4.4 Age verification

The stratigraphic units of each collection (e.g. formations) and their age were directly gathered from the publications and downloaded from PBBD. For the collections gathered from the literature, we first assessed whether a justification of the age assigned was provided in the corresponding publication. The following justification criteria were used: (a) the publication describes the geology of the locality; (b) the publication provides a reference in which the age of the locality was explained; or (c) the publication describes the use of specific method of age determination, e.g., isotope dating. If at least one criterion was fulfilled, the collection was marked as *valid* in the “age_evaluation” column of the collection dataset, otherwise, it was marked as *invalid*. To evaluate publications written in a language different from English, Spanish, German, French, Czech and Russian, we translated them using Google Translate (https://translate.google.com). Whenever the quality or formatting of the text in the publication did not allow using of Google Translate, it was marked as *unverifiable* in the “age_evaluation”. These records should be re-visited and evaluated in the future. Age justifications were not evaluated for the PBDB collections in the same manner given that this data was downloaded without the associated publications. However, we marked PBDB collections as *valid/invalid* based previous works that had critically evaluated the neoselachian records in PBDB (Pimiento et al., 2016; Pimiento & Clements, 2014). Additionally, comments made by enterers of the collections to PBDB were checked and if explicit doubts about an age associated with a collection was reported, the collection was also marked as *invalid* (e.g., collection PBDB_ 204174).

For both the collections gathered from the publications and downloaded from PBBD, the age associated with stratigraphic units was checked against the most recent literature to assess whether it was up to date. Whenever the age of a collection was updated, a reference of the publication used to make the update was added to the “comments_evaluation” column. If the age of a collection could not be validated due to a lack of information in the source publication, this was also noted in the “comments_evaluation” column.

The age of the occurrences was also evaluated. To do so, we identified taxa with exceptionally broad age-ranges (i.e., more than 60 Myr). Then, we checked outlier occurrence age ranges against primary literature (e.g., Cappetta, 2012). Whenever we found that an occurrence that was far outside the known age-range of a taxon we marked it as *invalid* in the “age_evaluation” column of the occurrence dataset. Otherwise, the “age_evaluation” entry of a given occurrence was marked as it was for its corresponding collection.

## 3. Data Records

### 3.1 Overview

The version of the FINS dataset presented here contains 5,972 collections (54.64% from the literature and 45.36% from the PBDB, Fig. 2a) and 30,100 occurrences (56.70% from the literature and 43.30% from the PBDB; Fig. 2b). Most collections extracted from literature reported occurrences based on isolated teeth (92.55%), followed by vertebrae (6.83%) and dental plates (5.42% of collections; Fig. 2c). Fossil type was not reported in only a minority of collections (2.64%, Fig. 2c). Note that information on fossil type was only extracted from the literature and not the PBDB (see Data collection). After data validation (see Data harmonization, curation and validation), the ages assigned to 274 collections (4.59%) were marked as *invalid* or *unverifiable.* From these, four were marked as invalid based on age-evaluations performed on previous studies (Pimiento et al., 2016; Pimiento & Clements, 2014). Furthermore, 1,424 occurrences (4.73% of total occurrences) were marked as *invalid* or *unverifiable,* out of which 1,101 were invalid on the basis of age (77.32%; Fig. 2d), 323 on the basis of taxonomy (22.68%; Fig. 2e), and 24 on the basis of both (1.69%). Of all occurrence names, 25.74% represented outdated synonyms (Fig. 2e). The majority of these synonyms (95.75%) were updated to currently valid names, while 329 of them represented dubious nomenclature. Considering the dubious nomenclature, a total of 184 names (88 from literature and 96 from PBDB) were marked as *unverifiable*. One occurrence from PBDB was marked as *nomen dubium* (n. gen. Agabelus n. sp. porcatus; Cope, 1875) and 144 occurrences from the literature were marked as *nomen nudum* based on the SR taxonomic database (Fig. 2e). Additional 17 occurrences were marked as invalid based on taxonomy after closer inspection of the source literature or based on previous studies(Pimiento et al., 2016; Pimiento & Clements, 2014). Considering taxonomic evidence, 18.44% of occurrences extracted from literature and 64.76% of occurrences downloaded from PBDB lacked evidence in the source publication, resulting in an overall 38.50% of the total occurrences lacking evidence (Fig. 2f). Overall, 2342 occurrences were reported with uncertain nomenclature (7.78% of total occurrences), of which 50.60% include “*cf”*, 22.89% “*aff”*, 18.49% “?”, and 9.48% are names with quotation marks.

**Figure 2.**
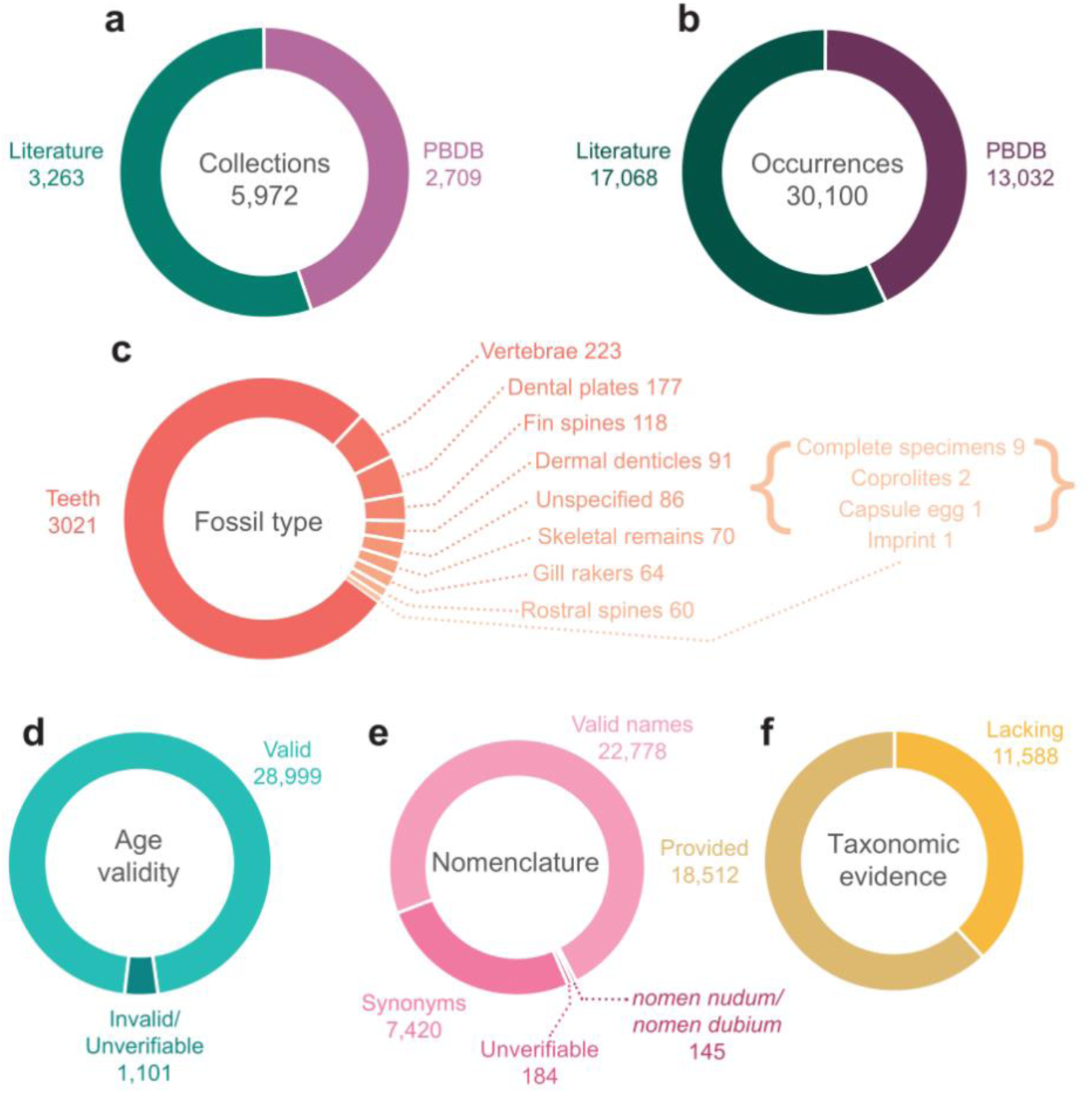
Overview of the FINS data. Total number of collections (a) and occurrences (b) present in the dataset and their proportion based on the source (literature and PBDB). c) Proportion of fossils types in the collections extracted from literature. d) Proportion of occurrences with age assignment marked as valid, invalid or unverifiable based on whether it was justified in the source publication (see 2.4. Data Evaluation Curation and Validation). e) Proportion of occurrence taxonomic assignments which were reported under a currently valid name in their source publication (*Valid name*), which were reported using an outdated synonym (*Synonyms*), which were reported under a name now considered a *nomen nudum* or *nomen dubium*, and which were not present in our taxonomic list and could not be evaluated (*Unverifiable*; see 2.4. Data Curation and Validation). f) Proportion of occurrence taxonomic assignment for which evidence was provided or lacking in the source literature.

### 3.2 Neoselachian literature

The data downloaded from PBDB is associated with 1,139 publications. We extracted data from 851 additional publications which we identified using SR. In total, FINS includes data extracted from 1,990 publications. The total number of neoselachian publications (from both PBDB and SR) has increased over time (Fig. 3A-B). The earliest publication reporting a fossil neoselachian occurrence entered in the PBDB is from 1839 on the genus *Squalus* from England (Bowerbank, 1839) (Fig. 3B). The number of publications on the neoselachian fossil record remained low until 1970. From 1970 to 2020, the numbers of publications per year showcase a steadily increasing trend, reaching the highest number in 2014 with 64 publications (Fig. 3A-B).

**Figure 3.**
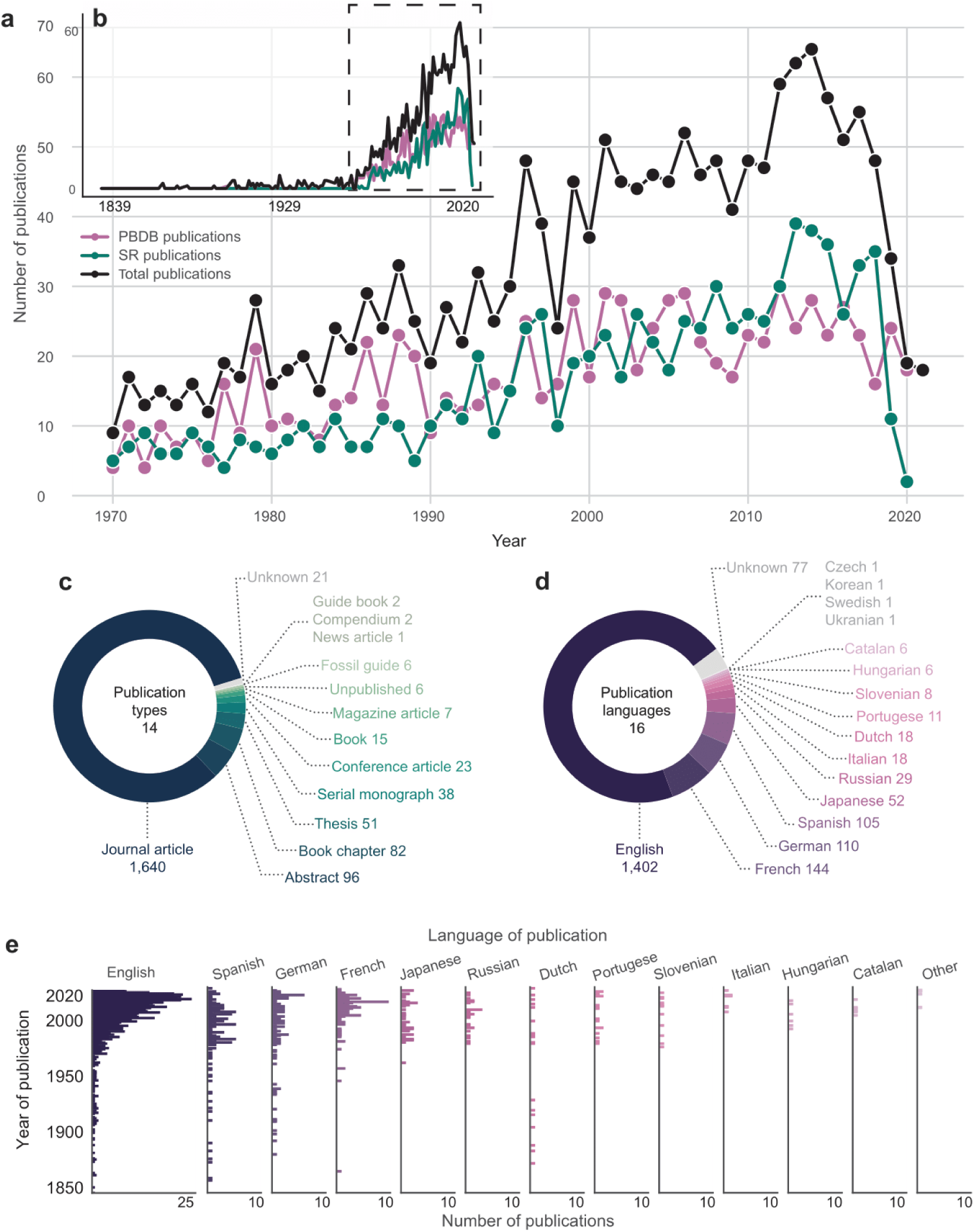
Overview of the literature on fossil neoselachians. a) Number of publications per year in total and split by the source [Paleobiology Database (PBDB) or Shark-References (SR)] published since 1970 until 2020; b) Number of publications per year in total and split by the source, spanning from the oldest publication entered in the PBDB (1839). Dashed-box denotes years 1970 to 2020; c) Types of publications gathered from both sources combined; d) Languages in which the literature was published from both sources combined; e) Number of publications per year by language, with the last plot combining the languages in which only one publication was published [Czech (2018), Korean (2018), Swedish (2003) and Ukrainian (2015)].

The majority of the publications assessed were peer-reviewed journal articles (82.41%), with the second most frequent publication type being abstracts (4.82%; Fig. 3C). Seven articles from non-academic magazines were also included in the data extraction process, as the source magazines were published by academic publishers and were focused on geology or palaeontology, for example the Dutch magazine *Grondboor & Hamer* published by the Dutch Geological Society.

Literature written in 16 languages was processed during the data collection. The most frequently published language is English (70.45% of all publications), followed by French, German and Spanish (7.24%, 5.53%, 5.28% respectively; Fig. 3D). Publications in English show steady increase over time, while being present in every decade since 1850s. Spanish, German and French publications also first appeared in the mid 1800s, and together with those in English, significantly increased from 1970 onwards. Although publications in Dutch also first appeared in the 1860s, they do not showcase the 1970s increase. The 1970s also mark the increase in frequency of publications in Japanese, Russian, Portuguese and Slovenian, and the beginning of publications in Italian, Hungarian and Catalan. The remaining four languages, Czech, Korean, Swedish and Ukrainian, are all represented by a single publication each, thus no trends can be observed (Fig. 3E).

### 3.3 Taxonomic composition

In total 2,014 unique taxa are present in the FINS dataset, including 507 taxa not yet entered in the PBDB. The taxa are composed of 1,607 unique species, of which 1,103 (68.64%) are selachimorphs (sharks), 502 (31.26%) batoids (rays and skates), and one additional species which is not resolved to the superorder level (*Nanocetorhinus tuberculatus*). There are 512 unique genera, of which 321 (62.57%) are selachimorphs, 190 (37.03%) batoids and one additional genus which is not resolved to a superorder level (*Nanocetorhinus*). Finally, there are 88 unique families [58 (65.91%) selachimorphs and 30 (34.09%) batoids] and 14 orders (10 selachimorphs and 4 batoids). The genera *Cretomanta*, *Odontorhytis* and their species are all considered to be selachimorphs, despite not having a fully resolved classification (Cappetta, 2012; Vullo et al., 2021). The most abundant species in the dataset is *Otodus megalodon*, represented by 606 occurrences (2.01% of total occurrences), followed by *Carcharodon hastalis* with 485 occurrences (1.61% of total occurrences). Regarding extant taxa, all 13 orders, 92.98% of families, 56.50% of genera and 12.07% of species are represented in the fossil record. The vast majority of occurrences in FINS are identified to the species-level (70.88%), with genus-level occurrences being the second most abundant (24.92%) and the remaining ranks representing only a small fraction (Fig. 4A). Selachimorphs have more occurrences than batoids (76.27% *vs* 22.45% of total occurrences; Fig. 4B). Lamniformes and Carcharhiniformes are the two most abundant selachimorph orders (37.45% and 21.53% of total occurrences, respectively) and Echinorhiniformes the least (0.35% of total occurrences). Myliobatiformes is the most abundant batoid order (12.29% of total occurrences) and Torpediniformes the least (0.29% of total occurrences; Fig. 4B).

**Figure 4.**
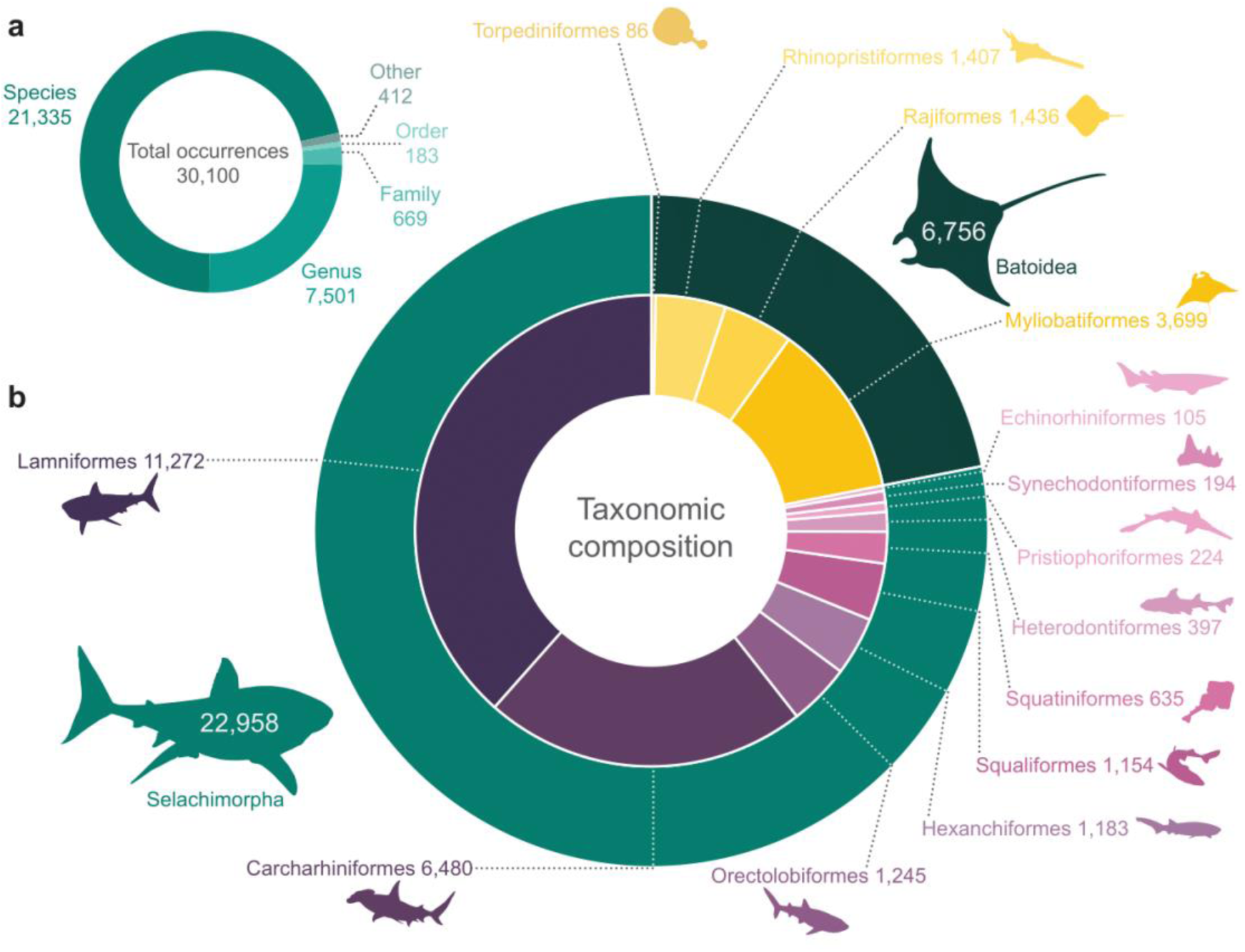
Overview of the taxonomic composition of the FINS dataset in terms of number of occurrences. a) Number of occurrences identified to different taxonomic ranks. b) Representation of Selachimorpha (sharks) and Batoidea (rays and skates) and their respective orders. Only taxa identified to at least the order-level were included. The Selachimorpha and Batoidea numbers are higher than just the sums of the numbers of orders, as these groups also include taxa which do not have a resolved order (see text). Outline images obtained from Phylopic (www.phylopic.org), Synechodontiform tooth outline adapted from Rees (2000), under CC BY 3.0. license.

### 3.4 Chronostratigraphic coverage

The majority of occurrences belong to collections with an age resolved to a stage (40.28%; e.g., Ypresian) or a sub-epoch (33.34%; e.g., middle Miocene). The mean age range of an occurrence is 7.7 Myr. Neoselachian fossils were found in every geological epoch from the Early Cretaceous to the Holocene, with Miocene, Late Cretaceous and Eocene showing the highest numbers of occurrences (26.70%, 26.68% and 20.43%, of all occurrences, respectively; Fig. 5A). All geological stages apart from Northgrippian (Holocene) were found to yield neoselachian fossils (Fig. 5B). Notably, the Holocene is the most fossil-poor epoch, yielding only 35 occurrences (0.12%), followed by the Pleistocene and the Early Cretaceous (1.69% and 3.68%, respectively).

**Figure 5.**
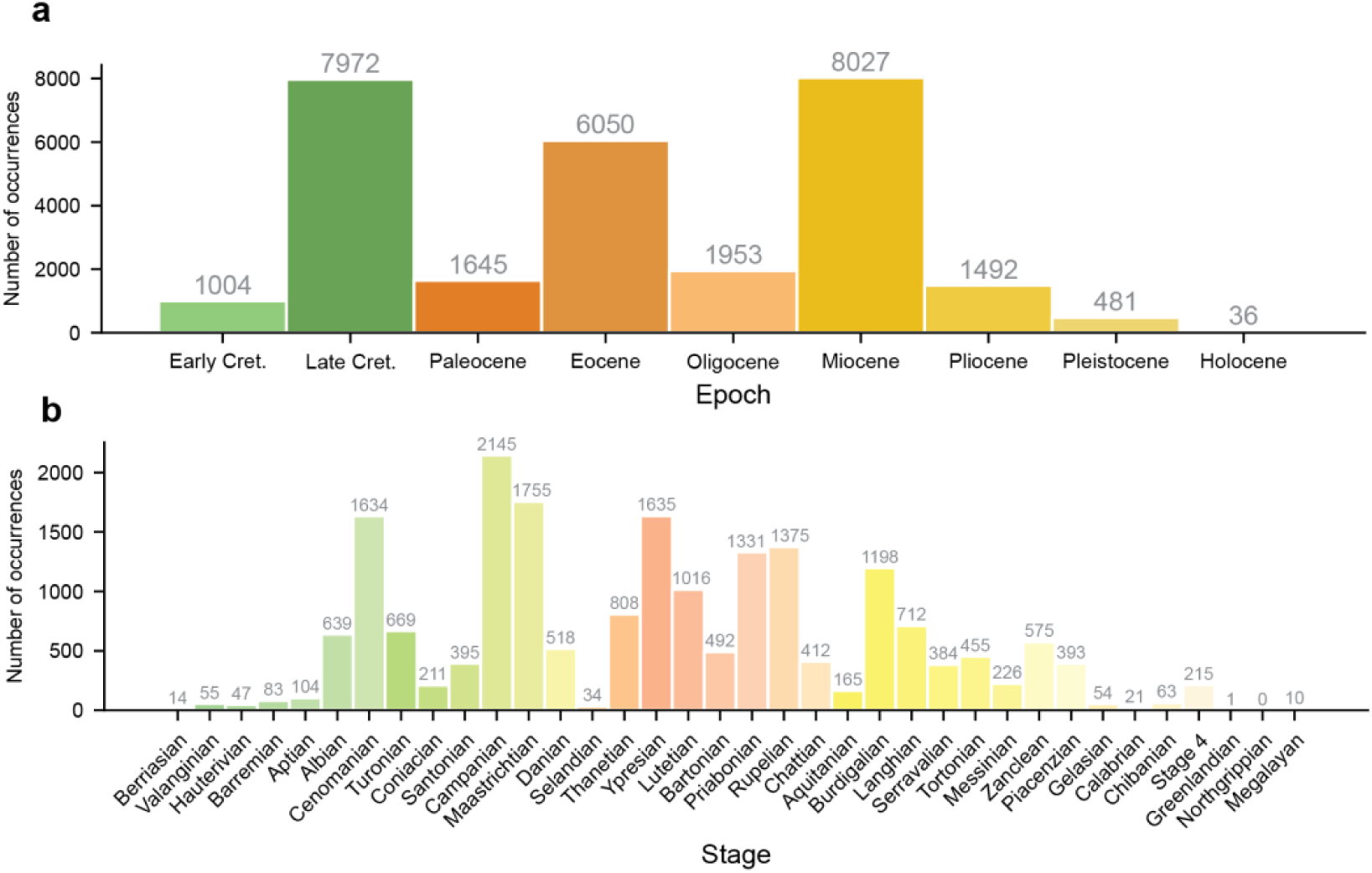
Abundance of neoselachian occurrences in FINS in the different geological a) epochs and b) stages.

### 3.5 Geographical distribution

Occurrences were identified from every continent, with Europe and North America being the most prolific, containing 41.75% and 27.90% of all occurrences, respectively (Fig. 6A). Indeed, the global North yields a much higher number of Neoselachian fossils relative to the global South (75.47% and 24.53% of occurrences respectively). Gaps in the geographical coverage of the neoselachian fossil record include East and Southeast Asia (with the exception of Japan); East, Southeast and Central Africa; the Guianas and Central South America; and Northeast Canada (Fig. 6A). Notwithstanding, only three collections reporting neoselachian fossils from China are included in the FINS dataset, with the SR database only containing four publications focusing on China from the Cretaceous and Paleogene, out of which three report only hybodonts (Gilles Cuny, 2017; Mo et al., 2016; Zhang, 2007). Compared with the data downloaded from the PBDB, the data gathered from the literature (based on SR) not only expanded the overall amount of data, but also the geographical coverage, especially in Europe, Asia, South America, and Antarctica, where PBDB occurrences constitute the minority (30.41%, 39.24%, 30.71%, and 33.49%, respectively; Fig. 6B). The opposite applies to Oceania, Africa, North America and samples from oceanic sediments, where higher proportion of the data comes from the PBDB (72.04%, 66.76%, 56.26%, and 84.88% respectively; Fig. 6B).

**Figure 6.**
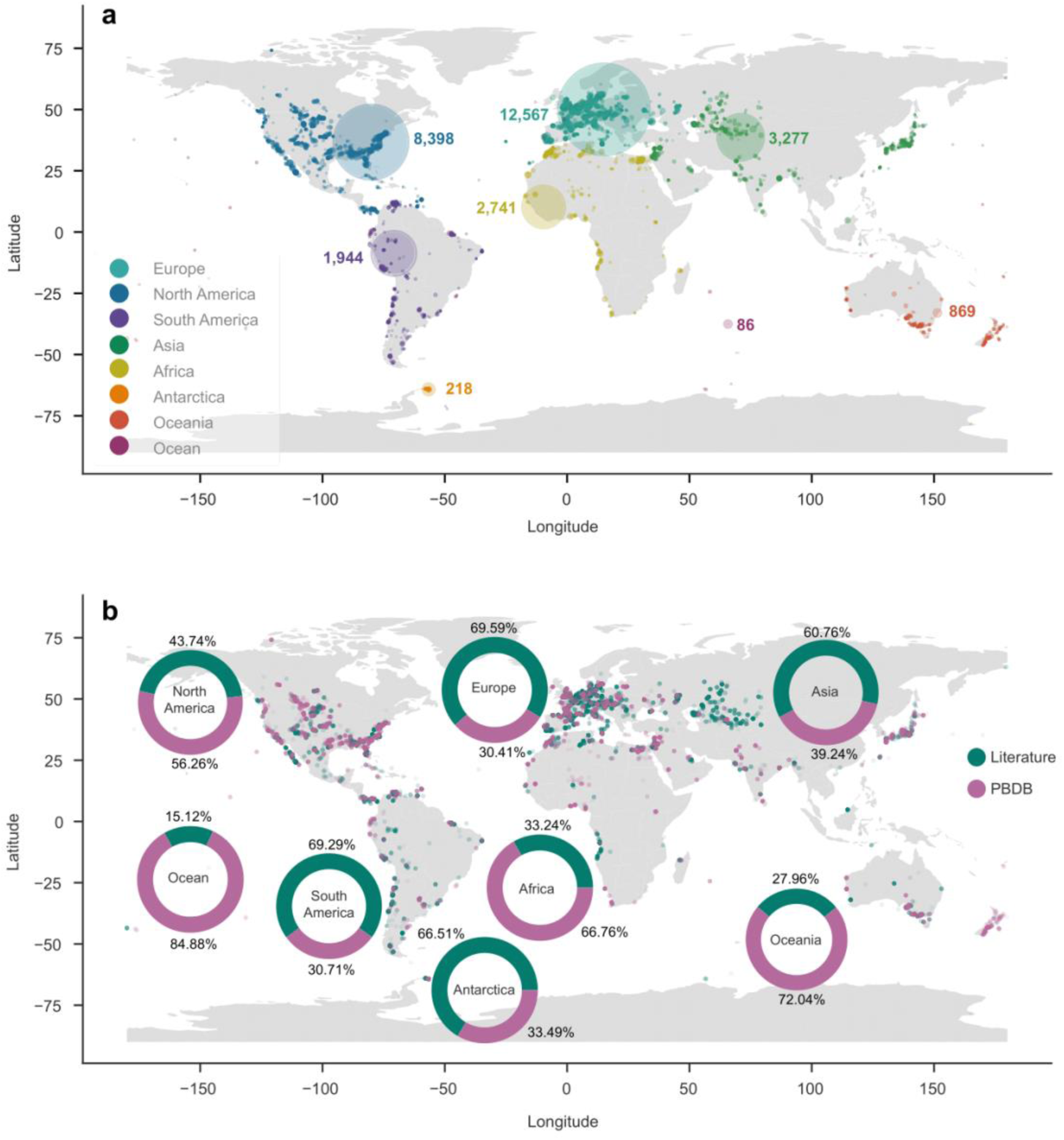
Geographical distribution of neoselachian occurrences in FINS. a) Global distribution of occurrences colour-coded according to the continent. Occurrences assigned to “Ocean” (presented in the Indian ocean) were reported from ocean drill cores. Large bubbles represent the proportional number of occurrences in the respective continent; b) Global distribution of occurrences colour-coded according to the source of the data – teal are data added to FINS from literature (from Shark-References), pink are data downloaded from the Paleobiology Database (PBDB), larger circles represent the proportion of data from PBDB and data extracted from literature.

### 3.6 Patterns and gaps

Overall, our synthesis of the literature demonstrates that the neoselachian fossil record is extensively documented in paleontological research. The earliest entry in the PBDB dates back to 1839, coinciding with the period of Agassiz’s landmark work on fossil fishes (Agassiz, 1833) which is considered the first comprehensive description of fossil sharks. This suggests that the PBDB effectively captures the beginning of our scientific understanding of fossil neoselachians. The number of publications has increased ever since, especially from 1970’s onwards (Figs. 3a,b) coinciding with the wider adoption of microfossil sampling techniques (Underwood, 2006).

The data collected in FINS further shows that the neoselachian fossil record is rich, with over 30,000 occurrences worldwide throughout the last 145 Myr (Fig. 2b). Notably, the vast majority of the fossil occurrences are represented by isolated teeth (Fig. 2c) and are identified to the lowest possible taxonomic unit: species-level (Fig. 4a). Indeed, the FINS dataset includes over 1,600 fossil species. Despite the fact that these taxa may not represent biological species (Wiley, 1978), they likely provide valuable insights into biological processes such as extinction and speciation. This is particularly important in paleobiology, as it has been argued that higher taxonomic ranks, such as genera or families, may not serve as accurate proxies for species-level diversity (Lloyd et al., 2012).

Our synthesis of the neoselachian fossil record further unveils some gaps in our knowledge. For instance, despite possessing the same “conveyor-belt” tooth replacement mechanism and shedding, batoids are significantly under-represented compared to selachimorphs. This likely stems from batoid teeth having single or double-layered enameloid as opposed to the triple-layered selachimorph teeth, which are likely to have higher preservation potential (Enault et al., 2013). Additionally, the collection method yielding the highest numbers of batoid teeth is screen-washing, whereas the most common collection method for neoselachian fossils is surface collection, favouring selachimorph teeth (Cappetta, 2012). A further bias towards larger specimens likely contributes to this pattern. The two most abundantly sampled species in our dataset are *Otodus megalodon* (606 occurrences), a species with the largest teeth found amongst sharks (e.g., Perez et al., 2021), and *Carcharodon hastalis* (485 occurrences), the teeth of which can reach relatively large sizes (e.g., Collareta et al., 2017). In contrast, shark species with smaller teeth, often associated with planktivorous diet, are much less abundant (e.g., *Megachasma applegatei* - 7 occurrences, *Aquilolamna milarcae* – 1 occurrence (Shimada et al., 2014; Vullo et al., 2021)). Additionally, larger teeth are often sought after commercially, which may increase the collection efforts focusing on such specimens, further shifting the bias towards larger species (Ragni et al., 2025). As many batoid species also possess reduced dentitions, tooth size may be a critical factor affecting their fossil representation. These factors combined likely contribute to the dominance of selachimorphs in the FINS dataset, with Lamniformes and Carcharhiniformes together representing 58.98% of total occurrences (Fig 4b).

The geological epochs yielding the highest number of neoselachian occurrences include the Late Cretaceous, Miocene and Eocene, together representing 76.89% of the total occurrences, whereas the most recent epochs (Holocene and Pleistocene) show the lowest number of fossil occurrences, with the Holocene being particularly fossil-poor. This finding is in line with the current knowledge of the neoselachian fossil record, showing that its quality decreases towards the present (Pimiento & Benton, 2020). The geographic distribution of the neoselachian fossil record appears to be skewed towards the global North, with notable gaps in Southeast Asia and Southeast Africa (Fig. 6). Interestingly, these regions are considered hotspots for modern elasmobranchs (Derrick et al., 2020; Lucifora et al., 2011; Pimiento et al., 2023). However, this observation is unlikely to reflect the true geographical distribution of fossils, and instead probably represents sampling effort, which is largely influenced by historical and socioeconomic factors (Raja et al., 2021). For example, even though South America represents only a small portion of all fossil occurrences in the FINS dataset, the number of publications focusing on marine fossils has been experiencing a boom in the last few decades (Chávez-Hoffmeister & Villafaña, 2023; Pimiento et al., 2013; e.g., Viglino et al., 2023; Villafaña & Rivadeneira, 2018), indicating a potential increase of described neoselachian fossils in the future. The scarcity of fossils from China is particularly puzzling, especially considering the region’s paleontological tradition (Zhou, 2022). Furthermore, the absence of any chondrichthyan fossil occurrences in the Geobiodiversity database (Fan et al., 2013), the largest database of fossils collected in China, further implies a significant gap in research on this group within the country.

## 5. Usage

Large-scale fossil occurrence datasets have played a pivotal role in advancing our understanding of the diversity of life on Earth by enabling a wide range of studies in palaeobiology (Allen et al., 2020; Boag et al., 2021; Chiarenza et al., 2023; Jones et al., 2022; Mathes et al., 2024). The FINS dataset was created with the aim to contribute to these fields by providing an openly accessible, comprehensive, and systematically curated global synthesis of neoselachian fossil occurrences from the Cretaceous to the Quaternary. With over 30,000 fossil occurrences spanning 145 million years and representing all major neoselachian clades, FINS offers an unprecedented opportunity to address an array of paleobiological questions in this major marine vertebrate group.

Two key assets of this dataset are its large number of occurrences and their fine taxonomic resolution. Multiple occurrences per taxon allow for more realistic speciation and extinction time estimates compared to those solely based on first and last appearances, as factors such as preservation can be taken in account (Silvestro, Salamin, et al., 2014; Silvestro, Schnitzler, et al., 2014). These improved estimates can in turn refine calibrations of molecular clocks and phylogenies and help identify times of elevated speciation and extinction rates. Similarly, large number of occurrences allows for more accurate reconstructions of past diversity patterns (e.g., R. B. Cooper et al., 2024). An occurrence-based dataset such as FINS can also be used to investigate the role of competition in diversification of neoselachian sub-groups (e.g., Condamine et al., 2019). Its fine-scale taxonomic resolution further allows these processes to be examined on a species level, potentially revealing patterns otherwise hidden when only higher taxonomic levels are considered. The same analyses can also be performed on a genus or family level to assess the robustness of the species-level results. Overall, the FINS dataset has the potential to provide insight into many crucial aspects of neoselachian evolutionary history.

The temporal coverage of the FINS dataset further offers the empirical foundation to explore how neoselachians responded to major global change such as those experienced across the K/Pg mass extinction, the PETM, or the Eocene–Oligocene cooling. Furthermore, it allows exploring how communities have responded to fluctuations in climate, sea level, and productivity across long timescales, shedding light on shifts in trophic structure, niche occupation, and ecosystem resilience. Furthermore, FINS’ broad geographic and stratigraphic coverage, as well as the detailed paleogeographic information associated to each occurrence, enables the reconstruction of past distributions, dispersal pathways, and biogeographic provinces, and allows testing palaeogeographical hypotheses about the timing and direction of range shifts, faunal interchange, and endemism. Together, these features make FINS a powerful and versatile resource for investigating the evolutionary and ecological history of sharks, rays, and skates through deep time.

Importantly, FINS was built with fully transparency, allowing users to tailor the data to the specific requirements of their analyses. Because each occurrence is associated with multiple categories of detailed information, the data can be filtered with precision. Users interested in a specific spatial region can utilise the categorical classifications of continents, latitudinal bands, or paleooceans, or define a specific area of interest using modern or paleo-coordinates. Those focusing on a specific subgroup of neoselachians can filter the data using multiple taxonomic categories, ranging from superorders to genera. Temporal windows can likewise be selected based on epochs, periods, or eras, or a specific time interval can be defined using the minimum and maximum ages. Data quality and reliability filters are also available through the taxonomy, age and evidence evaluation columns, as well as through a time interval category (i.e., stage, epoch, etc.) or by applying a specific age range cutoff value. All filtering categories can be combined, enabling users to create the exact subset of data needed for their question of intertest.

In addition to its detailed filtering options, the FINS dataset also allows the users to independently evaluate the data. Because taxonomic opinions evolve continuously, we retained the original taxonomic classifications exactly as given in the source materials. Access to the original data allows users to make an informed decision on whether to rely on the valid names assigned during the curation process (see 2.4 Data curation and validation), or to reassign names based on a different or more recent taxonomic frameworks. The inclusion of the original taxonomic classifications also enables the user to omit occurrences with uncertain classification (e.g., containing *aff.* or *cf.* classifiers) if desired. Users may also wish to consider how to handle duplicate records, defined here as multiple occurrences representing the same taxon in a single collection. Such cases may arise from two scenarios: i) two specimens from the same locality were assigned to a unique taxon in their source publication, however were deemed to be synonyms of the same taxon during the curation process (e.g., PBDB_24948 and PBDB_24947); or ii) a taxon was reported from the same locality in multiple publications (e.g., occurrences PBDB_12431 and PBDB_12438). If users prefer to remove such duplicates, this can be done by keeping only a unique combination of name and collection number. In sum, the data in FINS are presented transparently, giving users full flexibility to customise and critically assess the dataset to fit their research requirements.

## Supporting information

Data S1 - A list of synonyms and valid names

## Supporting Information

**Data S1.** List of synonyms and valid names from Shark References (https://shark-references.com). This list was used to assign valid names at the species- and genus-level to outdated synonyms.

## Data and Code Availability

The FINS dataset, version v1, is openly accessible at a Zenodo repository (https://doi.org/10.5281/zenodo.17371410). The codes used in Data merging and harmonization and Data curation and validation are provided in GitHub (https://github.com/Pimiento-Research-Group/FINS_dataset_harmonisation_and_curation), as well as the codes used in creating the figures presented in Data Records (https://github.com/Pimiento-Research-Group/Data_descriptor).

## Acknowledgements

Special thanks to J. Pollerspöck and N. Straube for providing PDFs from Shark References, which allowed us to extract most of the data in this dataset. We also thank them for providing their taxonomic list and for their comments and feedback during the preparation of this manuscript. We thank all the contributors who have entered data on neoselachian fossil occurrences into the PBDB, particularly M. Uhen, M. Clapham and M. Carrano, who entered the majority of these occurrences. We also thank the volunteers who helped along the long process of the dataset preparation and curation E. Ickin, S. Wüthrich, L.J. Guevara, A. Castro, L. Espinosa, and C. Jines. For their help with the translation of some French and Russian texts, we would like to thank V. Garnès, Vi. Šurmieliovas, and Va. Šurmieliovas. This project was funded by the PRIMA grant (no. 185798) of the Swiss National Science Foundation awarded to CP.

## Author contributions

CP conceived and coordinated the project. CP and JAV created the workflow. KK, JAV and AG collected data from literature. JAV curated the data obtained from PBDB. KK and curated the data from literature with the guidance of CP and JAV. AG and GHM, wrote and implemented the scripts for the estimation of paleocoordinates and paleooceans. KK led the writing with the guidance of CP and contributions from all authors.

